# Identification of watermelon genes involved in the ZYMV interaction through a miRNA bioinformatics analysis and characterization of *ATRIP* and *RBOHB*

**DOI:** 10.1101/2023.09.30.558769

**Authors:** Margarita Berbati, Maria Bousali, Athanasios Kaldis, Tomas Moravec, Timokratis Karamitros, Andreas Voloudakis

**Affiliations:** Laboratory of Plant Breeding and Biometry, Department of Crop Science, Agricultural University of Athens, 11855 Athens, Greece; Bioinformatics and Applied Genomics Unit, Department of Microbiology, Hellenic Pasteur Institute, 11521 Athens, Greece; Laboratory of Virology-Centre for Plant Virus Research, Institute of Experimental Botany of the Czech Academy of Sciences, Rozvojová 263, 165 02 Prague, Czech Republic; Laboratory of Medical Microbiology, Department of Microbiology, Hellenic Pasteur Institute, 11521Athens, Greece

**Keywords:** ALSV vectors, gene silencing, miRNAs, VIGS technology, ZYMV

## Abstract

Small RNA sequencing of healthy and ZYMV-infected watermelon (21 dpi) was performed and bioinformatics analysis identified 353 miRNAs from which 22 known and 331 new miRNAs. Important information about their precursors, their length, the loci of which they originated on watermelon genome are provided. The ZYMV genome could be a target for mir396a-3p, miR8706a, and miR1886i-5p from the miRbase, but none of them was identified in the watermelon miRNAome. Furthermore, watermelon miRNAome does not contain a miRNA targeting ZYMV genome with an expectation score ≤ 3.5. There were 34 resistance genes (CC-NBS-LRR, TIR-NBS-LRR, TIR-NBS) predicted as targets of 32 miRNAs in healthy watermelon. For nine differentially expressed miRNAs the respective target genes (10 in total) were bioinformatically predicted. For cla-new_miR307 (upregulated upon ZYMV infection) and cla-miR166h-3p (downregulated upon ZYMV infection) the targets were predicted to be *ClaATRIP* and *ClaRBOHB*, respectively, with *ClaATRIP* downregulated and *ClaRBOHB* upregulated upon ZYMV infection. ALSV-mediated VIGS of *ClaATRIP* rendered watermelon plants more resistant to ZYMV, whereas VIGS of *ClaRBOHB* resulted in higher levels of ZYMV titer in watermelon. These data suggest that *ClaATRIP* and *ClaRBOHB* are a susceptibility and resistant gene, respectively. Our results provide new insights in watermelon miRNAome and could propose new strategies for generating resistant watermelon to ZYMV.

**Highlights:**

- Bioinformatics analysis in healthy and ZYMV-infected watermelon plants gave 353 miRNAs, 22 known and 331 new.
- The watermelon miRNAome identified in the present study does not appear to target the ZYMV genome (sense and reverse complement).
- The differential expression of ten genes of watermelon in relation to ZYMV infection was validated.
- The silencing of the two genes, *ATRIP* and *RBOHB*, through VIGS strongly suggested that *ATRIP* is a gene of susceptibility and *RBOHB* is a gene of resistance.

## 1. Introduction

RNA silencing or RNA interference (RNAi) is a mechanism that plays an important role in plant protection against viruses. RNAi is induced after virus invasion in plant cells and its subsequent replication. Double-stranded RNA (dsRNA) is reproduced and recognized by plant dicer-like proteins (DCLs) which dice it in 21-25 nt short-interfering RNAs (siRNAs). One strand of the siRNA, designated the guide, is incorporated into the RNA-induced silencing complex (RISC) and directs RISC to the complementary mRNAs. A plant Argonaute protein (AGO), part of the RISC, slices the targeted viral RNAs reducing their accumulation and translation. The phenomenon is extended to the adjacent cells and in long distances through vascular system leading in systemic spread of RNAi. RNA-dependent RNA polymerase 6 (RDR6) subsequently acts on the cleavage products and converts them into dsRNAs for further processing [1–6].

Mature microRNAs (miRNAs) are 21 to 25-nt long molecules that regulate plant gene expression at the post-transcriptional level [7,8]. They are endogenous non-coding single-stranded molecules, originating from MIR genes and targeting endogenous mRNAs [9]. Mature miRNA duplexes are generated from primary miRNA transcripts (pri-miRNAs) with folding hairpin structure (stem-loop) containing base mismatches, loops, and bulges and are recognized by Dicer-like 1 (DCL1) proteins. The pri-miRNAs are processed in the nucleus to shorter RNA molecules of approximately 60 nts designated pre-miRNAs, which are exported in the cytoplasm and are further processed by DCLs to mature miRNAs. Plant miRNAs are involved in various biological activities, such as growth, development, hormone signal transduction, abiotic and biotic stresses [10–16].

Various silencing techniques have been developed, among them the virus-induced gene silencing (VIGS), a technique which employs recombinant viruses to specifically reduce endogenous gene activity (knockdown or knockout) and is based on post-transcriptional gene silencing (PTGS) resulting in the degradation of endogenous mRNA through a sequence homology-dependent RNA degradation mechanism [4,17]. VIGS is a suitable tool to study gene function and it has several advantages compared to other functional genomics approaches [18]. Plant gene fragments inserted in the virus genome become a source of siRNAs, which then target the corresponding plant gene mRNA for inhibiting its translation [19]. Thus, the expression of a specific gene can be downregulated through VIGS without the need to generate transgenic plants and the phenotype can be identified after the loss of the function for this specific gene within a single generation [20].

The VIGS vectors are binary Ti-plasmid derived vectors used for *Agrobacterium tumefaciens*-mediated plant transformation in which the viral genome is inserted. Fragments of 200 to 500 bp in length, targeting a complementary region of the target mRNAs are adequate. Plant inoculation with viral vectors is achieved via *A. tumefaciens* infiltration (agroinfiltration). The T-DNA containing the viral genome is then integrated into the host genome of at least one cell, transcribed and translated. This leads to the production of dsRNAs from which starts the RNAi [6,20,21]. Several virus vectors were designed for VIGS in plants and have been used successfully [19,22,23], e.g., tobacco mosaic virus (TMV), potato virus X (PVX), tomato golden mosaic virus (TGMV), and tomato yellow leaf curl China virus satellite DNA (TYLCCV), which have silenced the endogenous phytoene desaturase (*PDS*) gene in the model plant *Nicotiana benthamiana* [20]. The tobacco rattle virus (TRV) vectors have been developed for solanaceous plants [16,19] and the cucumber green mottle mosaic virus (CGMMV) vectors are appropriate for cucurbits [24]. There are also many other plant species, monocot and dicot, which have been silenced successfully using VIGS, such as *Hordeum vulgare, Zea mays, Oryza sativa, Triticum aestivum, Arabidopsis thaliana, Glycine max, Pisum sativum,* and *Malus domestica* among others [22,25]. One of the limitations of the VIGS technology is that the most reliable and effective VIGS vectors have a limited host range and are functional only in some plant species, mainly in the model plant *N. benthamiana* [26,27]. Moreover, some of the viruses used in VIGS induce symptoms that could challenge the interpretation of the phenotype caused by the silencing of the target gene [18].

Recently, an additional VIGS vector derived from the apple latent spherical virus (ALSV) was developed. ALSV is a small spherical virus which belongs to the genus *Cheravirus*, family Secoviridae, and has two segments of single-stranded, positive sense RNA with RNA1 encoding all proteins essential for genome replication and RNA2 encoding the movement protein and the three capsid proteins known as VP25, VP20, and VP24 [20,28–30]. It was isolated from an apple tree in Japan and, although it has only apple as a natural host, it has a wide range of experimental species including *Arabidopsis thaliana*, solanaceous, cucurbits, legumes and fruit trees [20,31]. Its advantage is the fact that it does not cause obvious symptoms in these hosts, although it infects them. This feature is particularly interesting and gives to researchers the ability to use it as vector for VIGS; it is reported that ALSV-based vectors effectively induced stable VIGS of endogenous genes for long periods in plants [20,29,31,32].

Zucchini yellow mosaic virus (ZYMV) is a single-stranded positive sense RNA filamentous virus, belonging to the genus *Potyvirus*, family Potyviridae, that causes serious damage in cucurbit crops worldwide [33]. The virus is transmitted by aphids in a non-persistent manner and the symptoms include severe mosaic, malformation of leaves and fruits, yellowing, stunting of the whole plant and of course, degradation of the product quality. It is present in most countries in the world including the Mediterranean countries (Italy, France, Cyprus, Algeria, Morocco, Israel, Greece etc.). In Greece, it is common in south regions since 1991 [34].

In order to limit the losses from ZYMV, our lab has employed several approaches to induce resistance against ZYMV infection in cucurbits based on exogenous application of dsRNAs [35] as well as amiRNAs [36]. In the present research, we performed NGS analysis in watermelon to identify the miRNAs that are differentially expressed due to ZYMV infection and to find their potential target host genes. Then, we employed VIGS in order to investigate the role of two host genes, namely *ATRIP* and *RBOHB* in the watermelon-ZYMV interaction. Our analysis indicated that *ATRIP* and *RBOHB* are susceptibility and resistance genes against ZYMV in watermelon, respectively.

## 2. Materials and methods

### 2.1. Plant growth conditions, ZYMV inoculum, and experimental workflow

*N. benthamiana* and watermelon [*Citrullus lanatus* (Thunb.) Matsum. & Nakai] cv Charleston gray were used. All plants were grown in a growth chamber under 25/22 °C (day/night) and 16 h day light conditions; plants were kept for observation for 30 days.

The inoculum for ZYMV-DE strain (DSMZ No. PV-0416) was obtained from the Leibniz-Institute DSMZ (*Deutsche Sammlung von Mikroorganismen und Zellkulturen* GmbH, Germany) as freeze-dried infected tissue and was propagated on zucchini plants (*Cucurbita pepo* var. Amerigo). Fresh leaf tissue with mosaic was collected 8 to 14 days after inoculation and was stored in −80 °C for using it as inoculum. Mechanical inoculations for ZYMV, using 20 μl sap, was carried out onto the adaxial surface of carborundum-dusted watermelon of the second true leaf. The sap was prepared from 1 g of well-homogenized tissue in 2 ml water [35].

The workflow followed in the present study for the validation of watermelon genes involved in the interaction with ZYMV, the VIGS of endogenous genes (namely *ClaATRIP* and *ClaRBOHB*), and the study of their effect in watermelon-ZYMV interaction is summarized in Fig. 1.

**Fig. 1.**
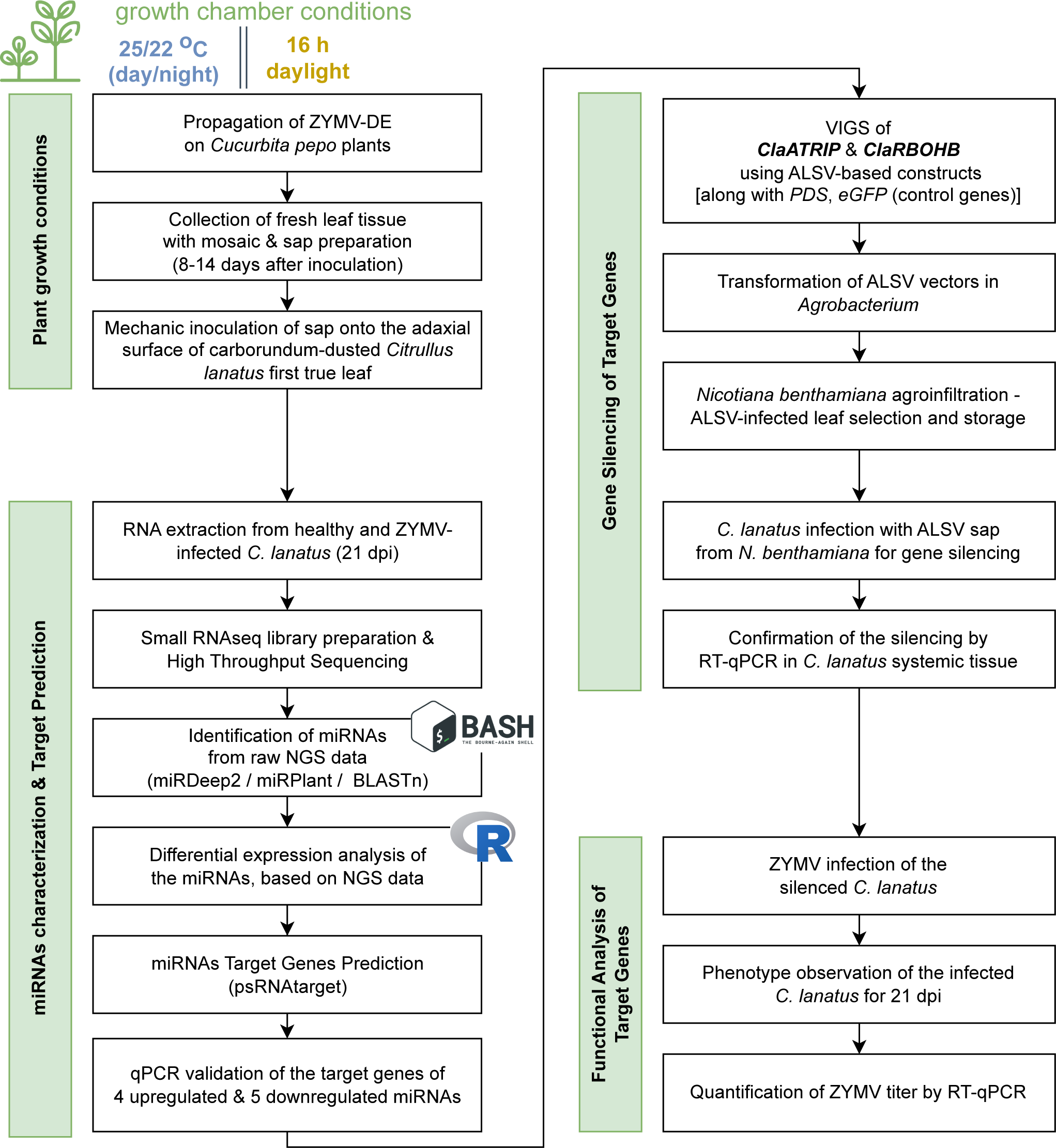
The workflow chart is presented the sequence of tasks performed during the experiments; miRNAs identification and characterization and the prediction, silencing and functional analysis of the target genes. Moreover, it gives information about the plant growth conditions, the origin of inoculum and the software tools to generate the list of miRNAs.

### 2.2. Total RNA isolation, RNAseq, bioinformatics identification of miRNAs, and their predicted mRNA targets

Total RNA isolation was carried out by TRIzol method according to the company’s instructions [35,37]. RNA samples from healthy and ZYMV-infected watermelon plants (21 dpi) were sent to Fasteris SA (Geneva, Switzerland) for high throughput sequencing, resulting in the generation of a small RNA library. Bioinformatics analysis of the NGS data involved the combined use of two software tools, miRDeep2 [38] and miRPlant [39], along with BLASTn [40] and in-house developed scripts written in R language [41]. In brief, Quality Control of the raw reads was conducted using FASTQC [42], followed by Adaptor Trimming using the CutAdapt v.1.15 tool [43]. Alignment of the reads to the *Citrullus lanatus subsp. vulgaris cv. 97103* Reference Genome [44,45] was performed with Bowtie v.1.2.3 [46], whilst the sequences of all hairpin and mature miRNA sequences that were used as input to the miRDeep2 and miRPlant pipelines, were accessed and downloaded from the miRBase v.21 [47,48]. We classified the identified miRNAs as: a) known or new, based on their similarity to previously reported miRNAs at the miRBase database, and b) up- or down-regulated, based on their expression levels in healthy and infected plants. For each miRNA, the formula log_2_(RPM from infected /RPM from healthy plants) (where RPM denotes reads per million) provides the measurement for the differential expression. The psRNATarget V2 (2017 update) (https://www.zhaolab.org/psRNATarget), was utilized to identify the potential targets of the miRNAs predicted, using the default cut-off parameters (maximum expectation: 5, number of mismatches allowed in seed region: 2, length of complementarity scoring [hsp]: 19, translation inhibition range 10-11 nt). All predicted target genes were blasted in the cucurbit genomics database (http://cucurbitgenomics.org/) to identify their location in the watermelon genome and their function. We selected five strongly downregulated and five strongly upregulated miRNAs for further study. The validation of their targets was performed by RT-PCR and RT-qPCR with specific primers (Table 1).

**Table 1.**
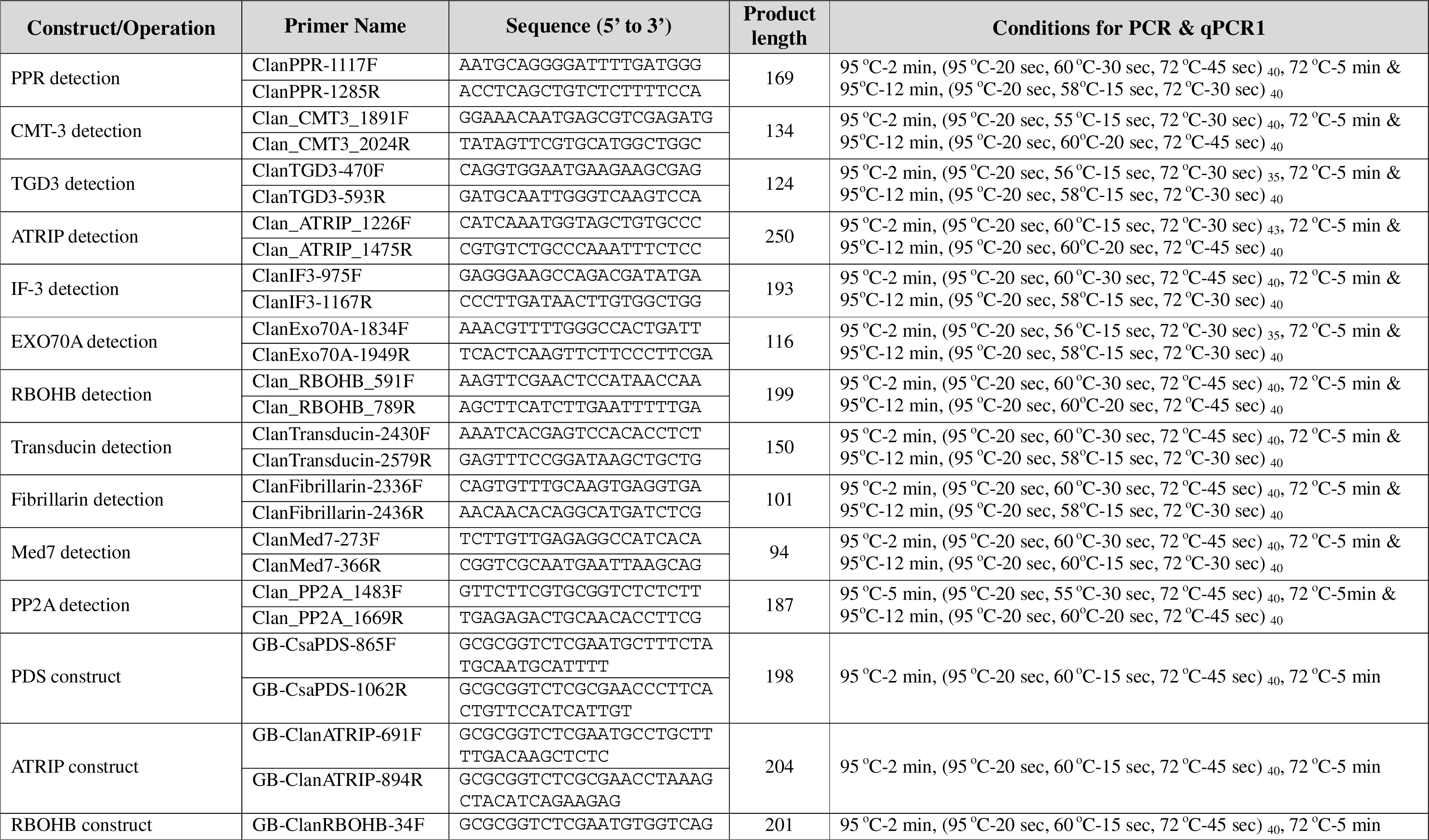
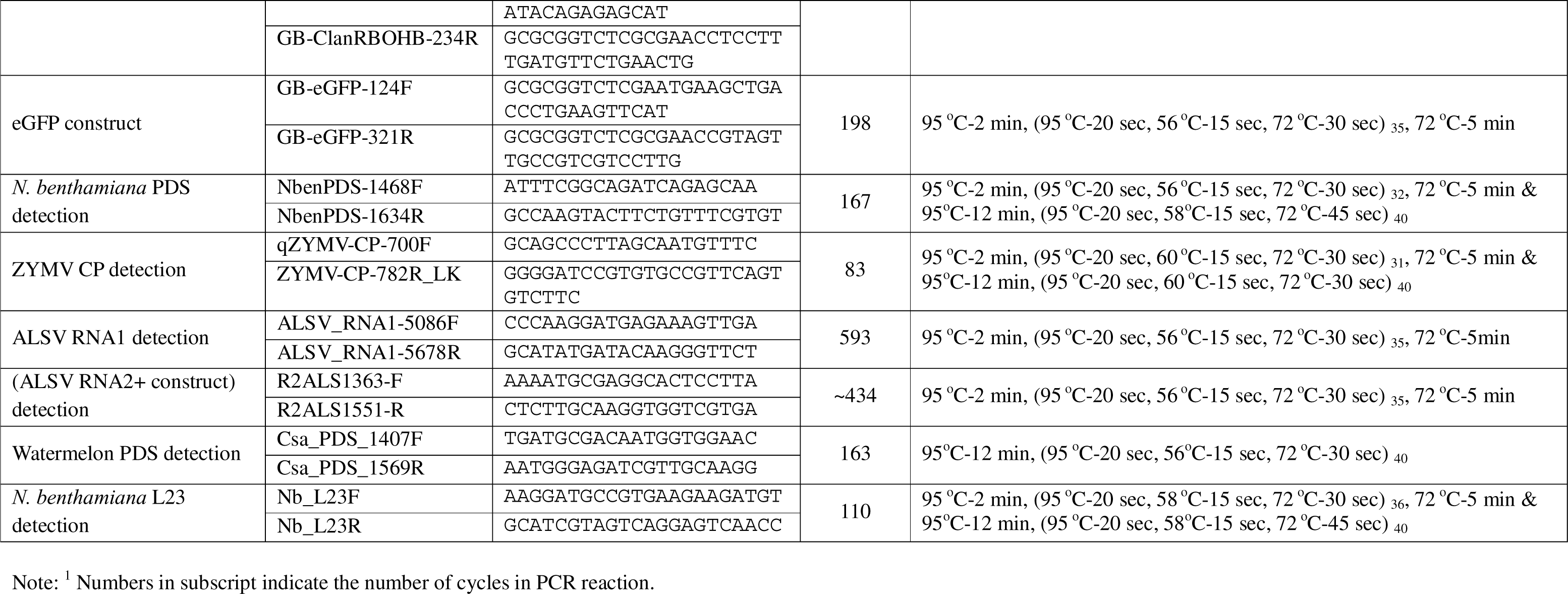
List of primers used in the present study and the conditions of PCR and qPCR.

We decided to study ten target genes of five downregulated and of four upregulated miRNAs in relation to ZYMV infection [11]; one upregulated miRNA had two potential gene targets. For the selection of the gene targets, we set the following criteria: a) the miRNAs that target them were strongly differentially expressed (differential expression value was higher to 2 or lower to −2); b) The expectation value given by psRNATarget for the target gene should be lower than 2. The lower the number the bigger the possibility was for the gene to constitute target of the miRNA; c) The bibliographic references of these targets should be related to RNA silencing or immunity of plants, i.e., for *RBOHB* there were references that it was involved to TMV resistance [49] and immunity in *N. benthamiana* [50] or nematode resistance in *Arabidopsis* [51], for *ATRIP* there is a proposal for its involvement in the programmed response of the plant to DNA damage [52]. Finally, ten target genes were selected and they are presented in Table 2.

**Table 2.**
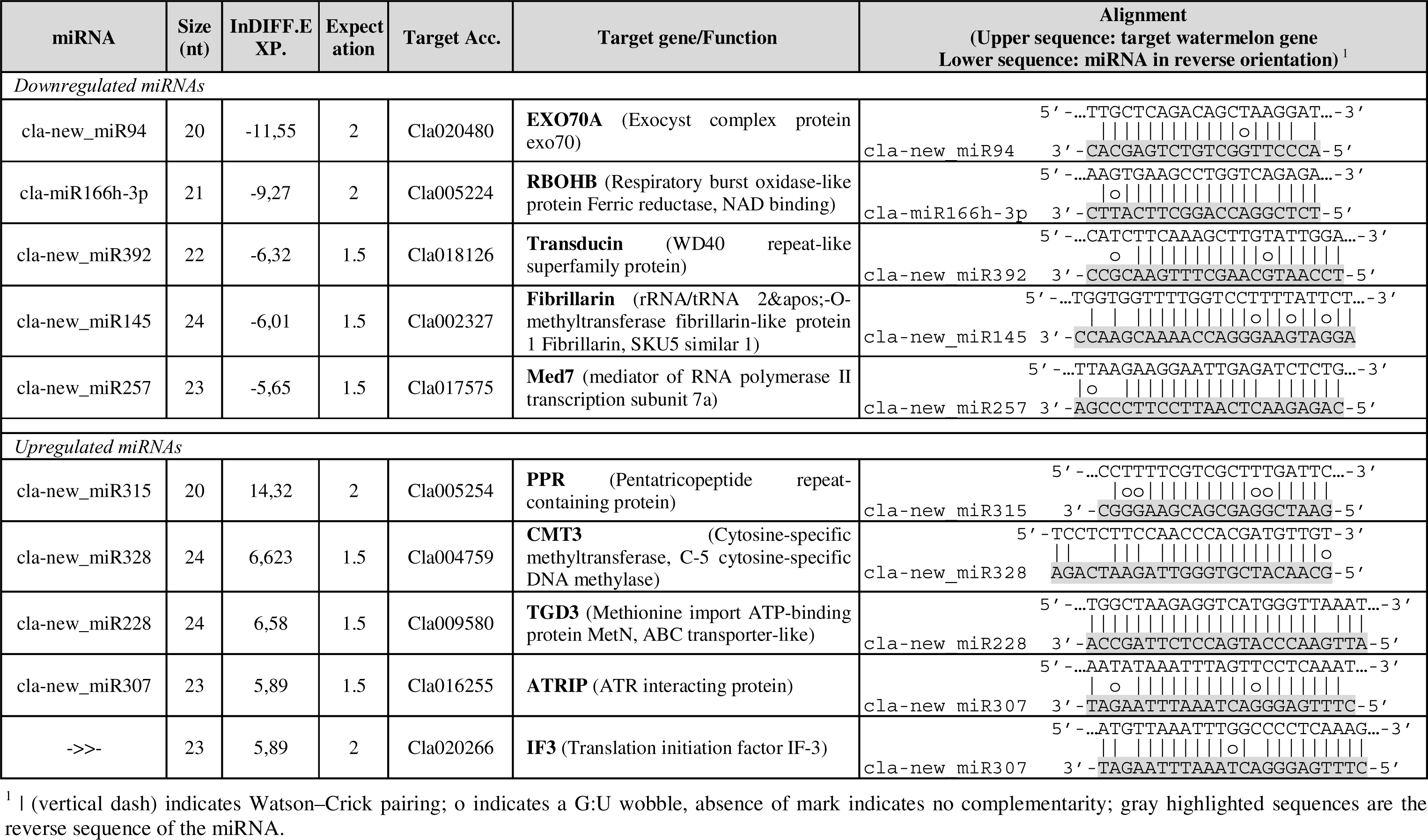
List of the selected miRNAs for their target validation. In list it is referred the name of the miRNA, the number of nucleotides, the expectation score, the accession number of the target gene and the name of target gene with some information of its function.

### 2.3. In silico detection of plant miRNAs targeting ZYMV genome

Plant miRNAs play a role in defence against virus infection in a process called miRNA-mediated resistance that could be manifested in a direct or an indirect manner. A bioinformatics approach [53] was followed to detect whether the ZYMV genome (sense and reverse complement) could be a direct target of miRNAs. The psRNATarget V2 was employed using ZYMV genome (OQ335839.1) as the target candidate, all the plant miRNAs present in the database were selected (accessed on September 2023), and the default parameters of the analysis tool as suggested.

The small RNA-target functional tab of psRNATarget V2 was employed to detect whether the watermelon miRNAome possess miRNA that is able to silence the ZYMV genome. The 353 watermelon miRNAs identified in the present study were used as small RNAs and the ZYMV genome (sense strand and reverse complement) was used as a target.

### 2.4. VIGS functional analysis of ClaATRIP and ClaRBOHB via ALSV-based constructs

Two of the putative target host genes in watermelon, namely the Respiratory Burst Oxidase Homolog B (*ClaRBOHB*) and the ATR interacting protein (*ClaATRIP*), were selected for VIGS in watermelon plants. By employing the GoldenBraid (GB) cloning method, we produced ALSV-based constructs for *ClaRBOHB* and *ClaATRIP* as well as for *ClaPDS*, and *eGFP* (enhanced green fluorescent protein gene) (the last two used as controls). In brief, for *ClaRBOHB*, *ClaATRIP*, *ClaPDS*, we performed Reverse Transcription (RT) of RNA isolated from healthy watermelon employing oligo-dT and random primers and PCR amplification employing specifically designed GB primers. Regarding *eGFP*, PCR amplification employing specifically designed GB primers was performed using as a template pBIN61-*eGFP* plasmid [54]. The PCR products were purified using a NucleoSpin Gel and PCR Clean-up kit (Macherey-Nagel, Germany); their concentration was brought to 30 ng/μl. Restriction-ligase reactions were performed in order to clone the PCR-amplified gene fragments onto the ALSV RNA2 genome. The components of the reactions were: pdb1a1 plasmid (80-100 ng), pUPD2 ALSV 5’ plasmid (80-100 ng), pUPD1 ALSV 3’ plasmid (80-100 ng), purified PCR product (50-80 ng), 1 μl of 10x T4 ligase buffer, 1 μl of T4 ligase (Promega, USA), and 1 μl of *Bsa*I enzyme (New England Biolabs, USA). The PCR program was: 30 min at 37 °C, 20 cycles of 2 min at 37 °C and 5 min at 16 °C and finally, 20 min at 37 °C and 20 min at 80 °C. The pdb1a1 plasmid is a GoldenBraid [23] binary destination vector. The pUPD2 ALSV −5’ plasmid is the part with 35S promoter and 5’-part of ALSV RNA2. The pUPD1 ALSV 3’ plasmid contains the 3’-part of ALSV RNA2 and the NOS terminator.

*E. coli* stellar cells (Takara, JP) were used for transformation by the heat shock method with well-established protocol. Transformed *E. coli* cells were selected in LB agar plates supplemented with kanamycin 50 μg/ml. Colony lysis PCR was performed for cloning confirmation purposes. Two single colonies (producing the desired PCR product) were selected for plasmid mini preps, performed with the NucleoSpin Plasmid kit (Macherey-Nagel, Germany); the successful cloning was verified by Sanger sequencing.

### 2.5. Agroinfiltration and ZYMV inoculation process

Each cloned plasmid was transformed to GV3101 *Agrobacterium* competent cells employing the freeze-thaw method [55]. Agrobacteria were plated on LB agar plates with antibiotics (Rifampicin 50 μg/ml, Gentamycin 100 μg/ml, and Kanamycin 100 μg/ml) and kept at 28 °C for 4 days. Colony lysis PCR was performed on single colonies to confirm the successful transformation; transformed Agrobacteria were stored at −80 °C. The agroinfiltration mixture for each gene (*ClaPDS*, *ClaATRIP*, *ClaRBOHB*, and *eGFP*) contained GV3101 cells harboring the corresponding gene fragment incorporated in the ALSV RNA2, EHA105 cells harboring the ALSV RNA1 [23], and GV3101 cells harboring the P19 silencing suppressor [56], in a 1:1:1 ratio and final OD_600_ of 1. The mixtures were agroinfiltrated in the abaxial surface of the third true leaf of *N. benthamiana* plants (3-week-old). At 10-14 days post agroinfiltration (dpa), top leaves were collected in order to be used as sap for mechanical inoculation of watermelon plants. Watermelon plants at their first leaf stage were used to be infected by ALSV using inoculum from ALSV-infected *N. benthamiana* leaves at their adaxial surface that was carborundum dusted. At 10-12 days post inoculation (dpi), systemic leaf samples (third leaf) were examined for confirming the silencing effect. The silenced watermelon plants were subsequently mechanically infected by ZYMV at the second leaf as in Kaldis *et al.* (1998). Three days post ZYMV challenge, samples from the fourth leaf were taken to determine the ZYMV titer. The silenced and ZYMV-infected watermelon plants remained at the growth chamber for 30 days for observation of the phenotype (symptoms).

We performed three experiments (biological replications) for *ClaRBOHB*- and *ClaATRIP*-silenced plants. Each experiment had the mock and the ALSV::*eGFP* as negative controls; each treatment contained 8-10 watermelon plants.

### 2.6. Plant RNA extraction, RT-PCR, and qPCR

RNA extraction was carried out by TRI reagent (MRC Inc, USA) method and reverse transcription was performed with FIREScript RT kit (Solis Biodyne, Estonia) with oligo-dT and random primers at 27 °C for 10 min, 37 °C for 60 min, and 85 °C for 5 min. For PCRs we used Taq polymerase (KAPA Biosystems, Cape Town, South Africa) and for qPCRs the FIREPol EVAGreen kit (Solis Biodyne, Estonia). The conditions for PCRs and qPCRs are shown in Table 1.

## 3. Results

### 3.1. Bioinformatics analysis of high-throughput sequencing of small RNAs

The NGS analysis of the small RNAs from healthy and ZYMV-infected watermelon plants identified 353 miRNAs, which mapped to the watermelon genome. The bioinformatics analysis detected their precursors which corresponded to pre-miRNAs and determined: a) the number of reads for each miRNA in the total reads (score) after normalization (reads per million, RPM), b) the number of the nucleotides of the mature miRNA, c) the chromosomal location for each miRNA, and d) their origin from the positive or negative strand (Table S1, Table S2, Table S3). In supplementary tables S1, S2, S4, and S5 the miRNAs are listed alphabetically.

197 out of 353 identified miRNAs were unique in healthy watermelon, 107 were unique in ZYMV-infected watermelon and 49 were common in healthy and ZYMV-infected (Figure 2A). 22 miRNAs were known and 331 were new (Figure 2A). The most numerous miRNAs were those of 23- and 24-nt-long that, interestingly, all belonged to the newly discovered miRNAs. For the 23-nt-long miRNAs they were 101 and 44 for healthy and ZYMV-infected watermelon, respectively, whereas for the 24-nt-long miRNAs they were 73 and 32 for healthy and ZYMV-infected watermelon, respectively (Figure 2). 309 of the mature miRNAs had only one precursor and 44 had more than one precursor. Differential expression analysis identified the upregulated and downregulated miRNAs as a result of the ZYMV infection. We identified 112 upregulated miRNAs in ZYMV-infected watermelon from which 107 unique and five common miRNAs (Figure 2A, Table S3). 214 miRNAs (197 unique + 17 common) that were expressed in healthy watermelon, were downregulated upon ZYMV infection (Figure 2A, Table S3). All miRNAs were deposited to the miRBase (https://mirbase.org).

**Fig. 2.**
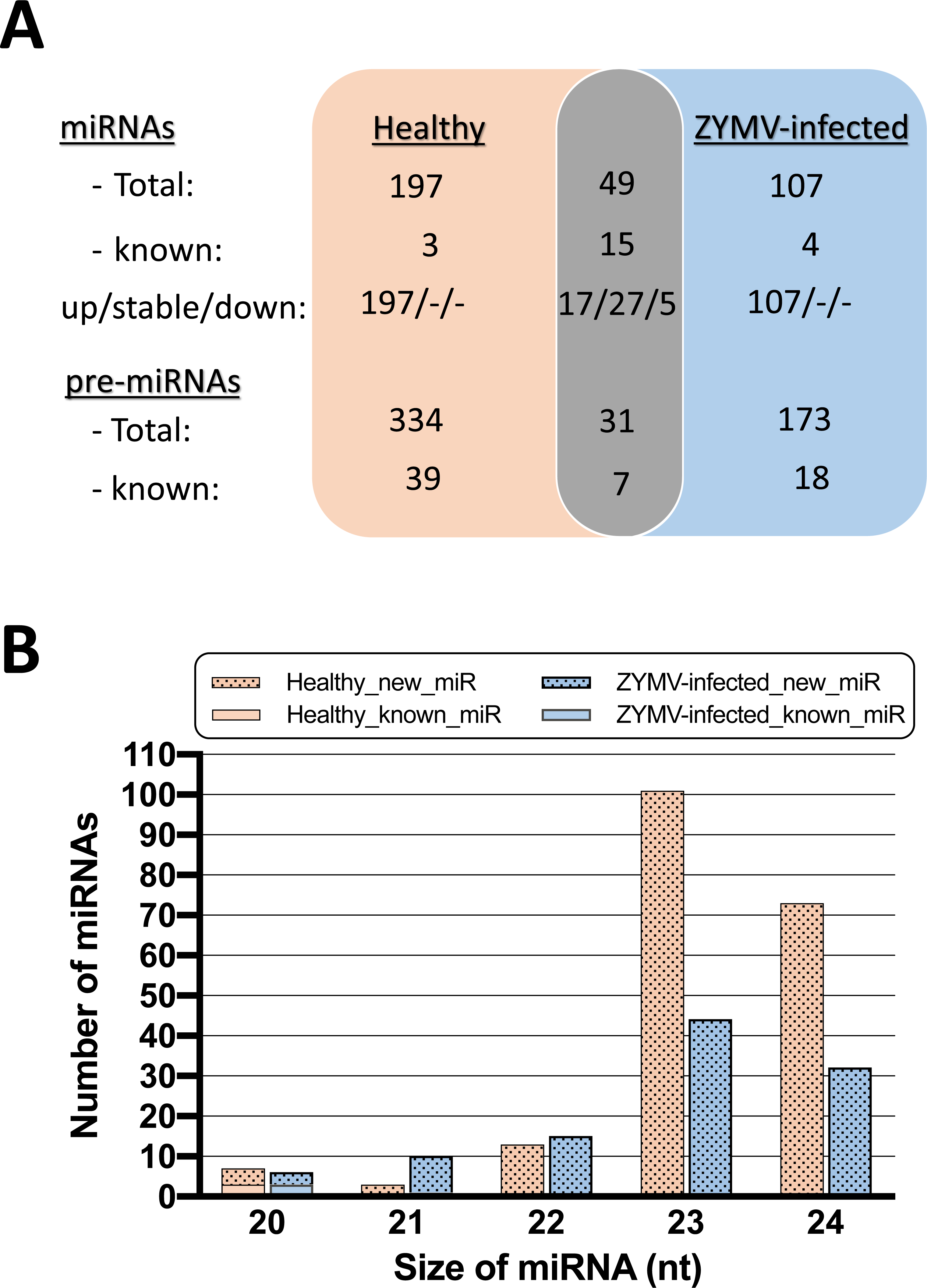
Watermelon miRNAome. A. Venn diagram showing number unique miRNAs and pre-miRNAs in healthy (left, light orange area) and ZYMV-infected (right, blue area) watermelon leaf tissue. The number of up-regulated, stable, and down-regulated miRNAs are also indicated. The grey highlighted area indicates the respective numbers for the common miRNAs. For the common miRNAs the number of up-regulated, stable, and down-regulated miRNAs are indicated in regards to the healthy tissue. B. Size classification of the unique watermelon miRNAs (20, 21, 22, 23, and 24-nt miRNA classes) shown in a stacked bar graph. Watermelon miRNAs in healthy plant (light orange) and ZYMV-infected (blue). For each miRNA size class, the non-dotted part of the columns (bottom) indicates the number of known miRNAs, whereas the dotted part of the columns (top) indicates the new miRNAs.

Using psRNATarget, the potential targets for each miRNA were obtained and found to be mapped to a wide range of loci on the watermelon genome. Each miRNA had more than one targets with a high or low expectation score (Table S4, Table S5). A number of 10397 and 7808 genomic positions were identified to be targeted by the miRNAs in healthy and ZYMV-infected watermelon, respectively (Table S4, Table S5). All predicted targets were blasted in the cucurbit genomics database (http://cucurbitgenomics.org/) to identify their location in the watermelon genome (column M in Table S4, Table S5) and their function (column O in Table S4, Table S5). There were 34 resistance genes (CC-NBS-LRR, TIR-NBS-LRR, TIR-NBS) predicted as targets of 32 miRNAs in healthy watermelon.

### 3.2. In silico detection of plant miRNAs targeting ZYMV

PsRNATarget output indicated that the miR396a-3p, miR8706a, and miR1886i-5p had an expectation score of 2.0 and thus having the possibility to post-transcriptionally silence ZYMV. All the above miRNAs are predicted to regulate the ZYMV RNA at the cleavage step. These three miRNAs were not present in the watermelon miRNAome, identified in the present study.

PsRNATarget using the watermelon miRNAome (353 miRNAs) to target the ZYMV genome (sense strand and reverse complement) resulted in no miRNA having an expectation score ≤ 3.5.

### 3.3. Validation of target gene expression for differential expressed miRNAs

RT-qPCR for *ClaEXO70A*, *ClaRBOHB*, *ClaTransducin*, and *ClaMed 7* confirmed that these genes were strongly upregulated (p<0.05), while *ClaFibrillarin* was slightly upregulated (Figure 3A). Furthermore, *ClaATRIP* was strongly downregulated as statistical analysis indicated at p<0.05; *ClaCMT-3*, *ClaTGD3*, *ClaIF-3*, and *ClaPPR* were slightly downregulated (Figure 3B). *ClaATRIP* and *ClaRBOHB* were selected for determining their involvement in watermelon-ZYMV interaction.

**Fig. 3.**
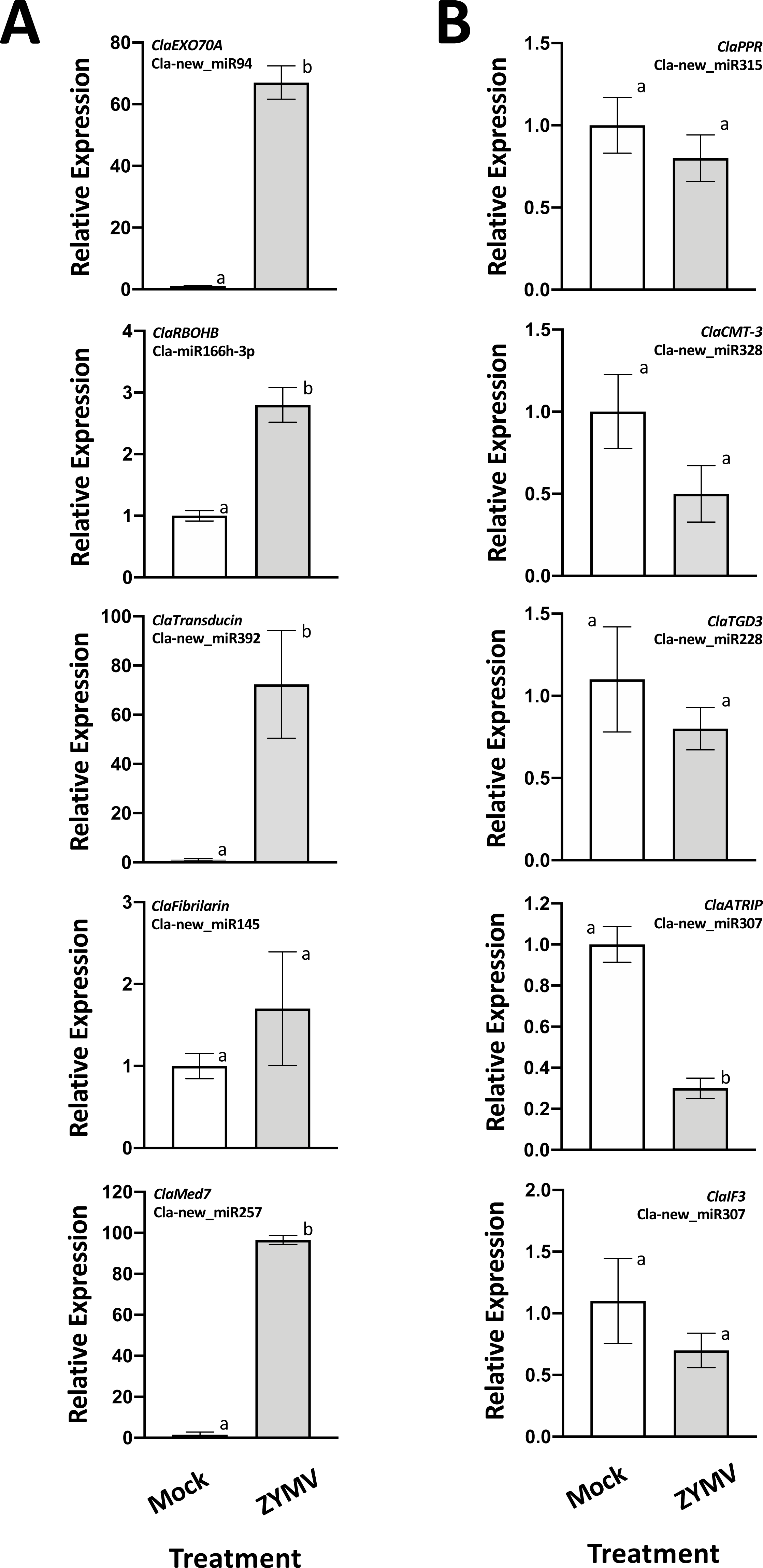
Quantification of the five A. upregulated and B. downregulated target genes in mock and ZYMV infected watermelon plants at 21 dpi and analysis by Schmittgen-Livak method. Statistical comparison by T-test for significance at p<0.05. White column: mock (healthy); grey column: ZYMV (infected).

### 3.4. Virus induced gene silencing (VIGS) in watermelon via ALSV

To determine the use of the ASLV as a VIGS vector in watermelon, we used it to silence the *PDS* (phytoene desaturase) gene. PDS is an enzyme which is involved in carotenoid biosynthesis pathway and its lack is responsible for the leaf photobleaching, a symptom which makes silencing visible (Figure 4A and Figure S1A) [19,29]. Since the percentage of infectivity (via agroinfiltration) of watermelon by ALSV was not reaching 100 % we decided to perform the infection of watermelon using *N. benthamiana* ALSV-infected sap (two-step procedure). The first step included the agroinfiltration of the ALSV-based construct with *PDS* in *N. benthamiana* (2-week-old) for virus multiplication and at 12 to 14 dpa *N. benthamiana* plants had their top leaves white (Figure S1A) and confirmed the gene silencing by RT-PCR and RT-qPCR (Figure S1B and S1C, respectively). The silenced *N. benthamiana* leaves were collected and used in the second step for mechanical inoculation of watermelon first true leaf [23]. Ten to twelve dpi the top watermelon leaves exhibited intense yellowing (Figure 4A), reminiscent of *PDS* silencing, with RT-PCR (Figure 4B) and RT-qPCR (Figure 4C) confirming it.

**Fig. 4.**
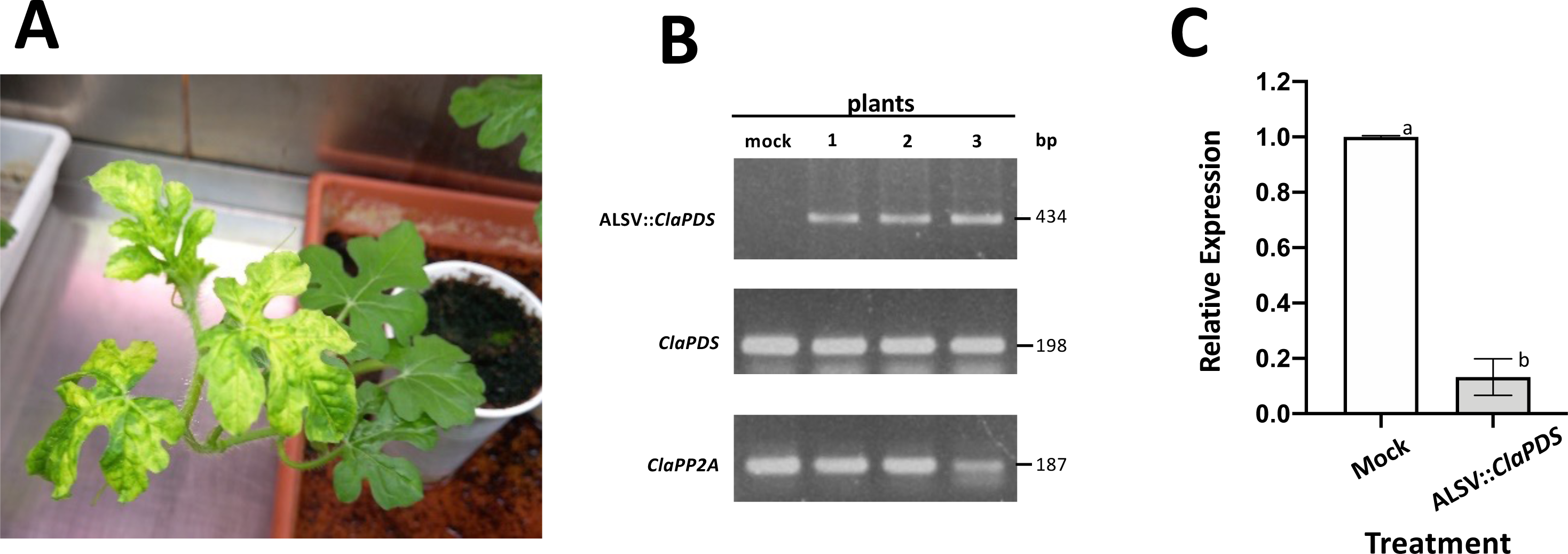
Virus-induced gene silencing of *ClaPDS* employing an ALSV-based construct in watermelon. A. Watermelon with white top leaves at 8 dpi with *N. benthamiana* extract (white leaves). B. PCR for mock and three watermelon plants at 8 dpi for ALSV::*ClaPDS* (construct), endogenous *PDS* and *PP2A* (control) gene. C. Quantification of *PDS* at 8 dpi. Quantification analysis performed by Schmittgen-Livak method. Statistical comparison of means done by T-test for significance, p<0.05. White column: mock; grey column: VIGS of PDS using ALSV::*ClaPDS*.

Subsequently, VIGS for *ClaATRIP* and *ClaRBOHB* was performed in watermelon. ALSV::*eGFP* was used as negative control. All watermelon plant groups were sap-inoculated with the ALSV constructs at the first true leaf. At 10 dpi watermelon plants showed a small reduction in size, due to ALSV infection, with no other visible symptoms (Figure 5A and 5C); samples were collected from the third leaf (Figure 5A and 5C) to validate *in planta* VIGS by RT-qPCR. In all experiments, we observed knockdown of *ClaATRIP* and *ClaRBOHB* expression as compared to the control group (plants treated with ALSV::*eGFP* construct) (Figure 5B and 5D).

**Fig. 5.**
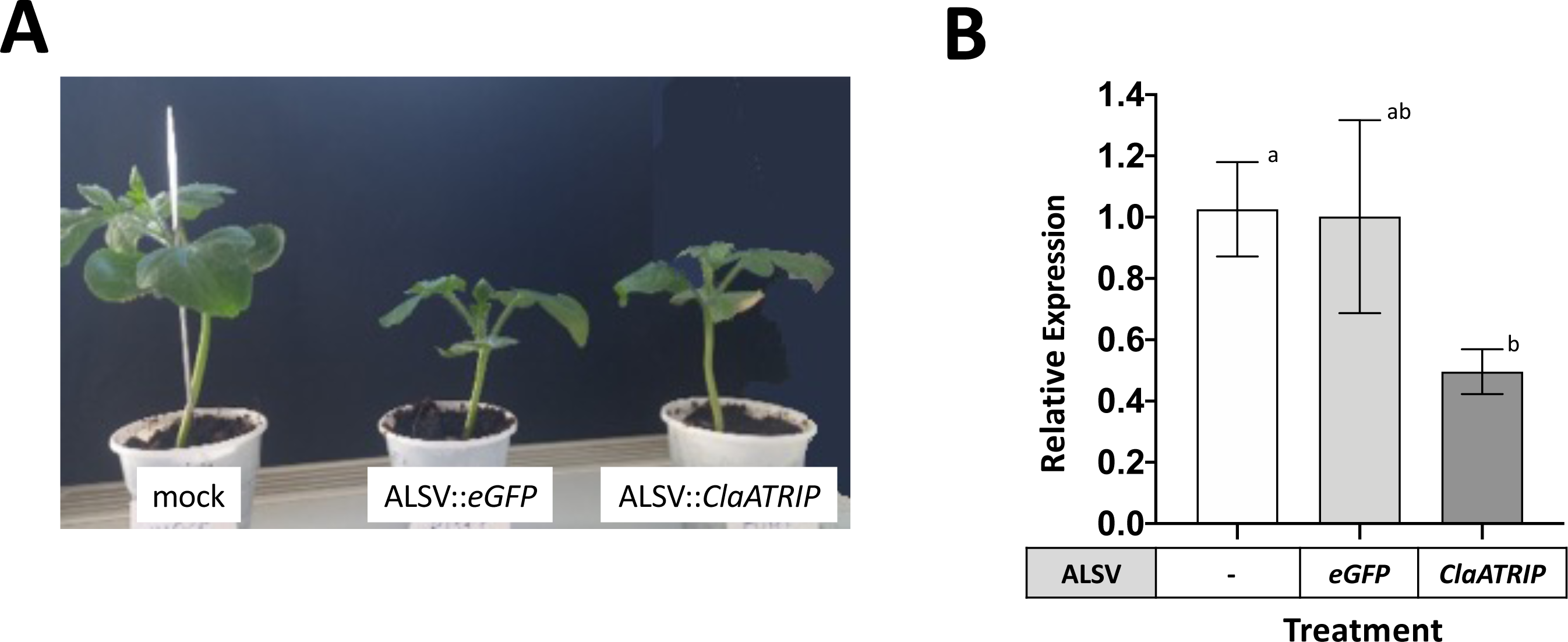
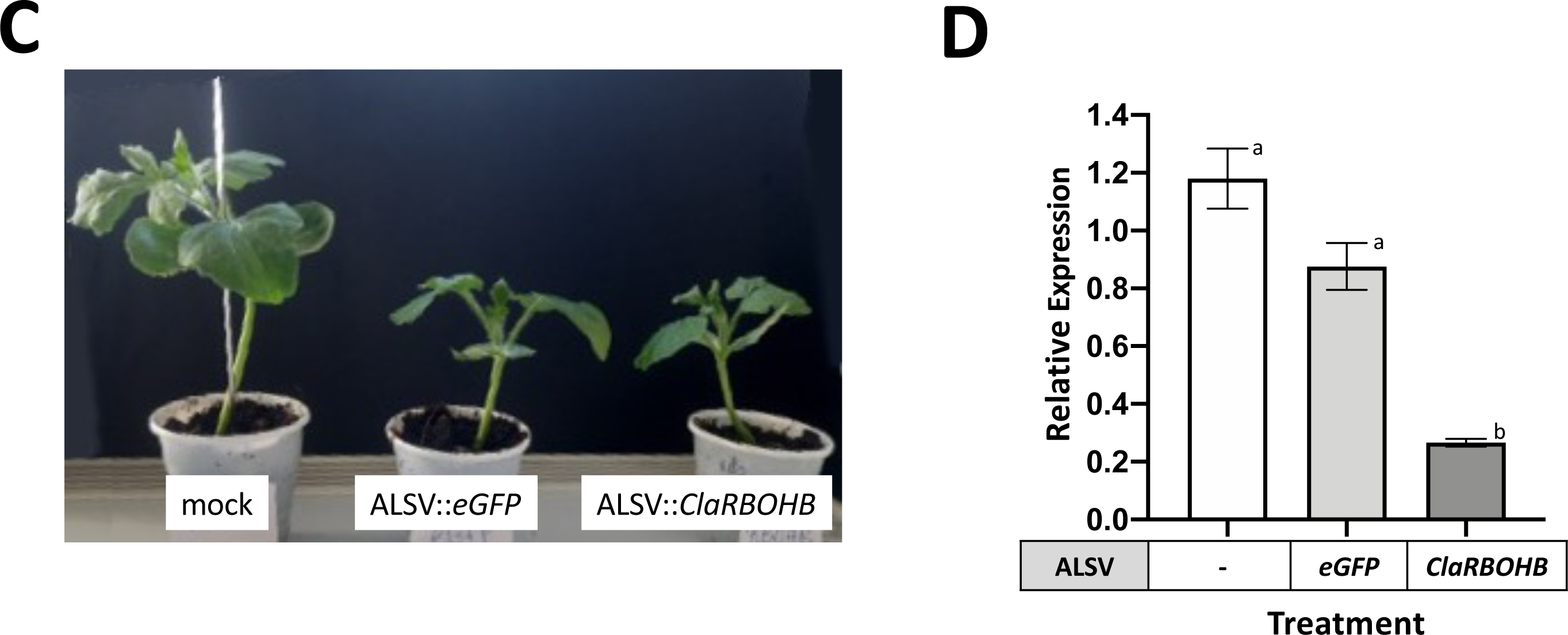
Virus-induced gene silencing of *ClaATRIP* and *ClaRBOHB* by employing ALSV-based constructs. A. Watermelon plants inoculated by *N. benthamiana* sap containing ALSV-based constructs at 10 dpi. Left plant: mock (not treated); middle plant: ALSV::*eGFP* inoculated; right plant: ALSV::*ClaATRIP* inoculated. B. Quantification of the *ClaATRIP* gene in watermelon at 10 dpi for silencing confirmation. C. Watermelon plants inoculated by *N. benthamiana* sap containing ALSV-based constructs at 10 dpi. Left plant: mock (not treated); middle plant: ALSV::*eGFP* inoculated; right plant: ALSV::*ClaRBOHB* inoculated. D. Quantification of the *ClaRBOHB* gene in watermelon at 10 dpi for silencing confirmation. Quantification analysis performed by Schmittgen-Livak method. Statistical comparison of means done by T-test for significance, p<0.05. White column: mock; grey column: *eGFP*; dark-grey column: *ClaATRIP*.

### 3.5. ZYMV infection of ATRIP- and RBOHB-silenced watermelon plants

The *ClaATRIP*- and *ClaRBOHB*-silenced watermelons were inoculated with ZYMV at the second leaf (12 dpa) and three days later samples from the fourth leaf were sampled to inspect the course of ZYMV infection. We performed RT-qPCR for ZYMV-CP to estimate ZYMV titer with the results presented in Figure 6. ZYMV-CP levels were low in *ClaATRIP*-silenced watermelon plants (Figure 6A) suggesting that *ClaATRIP* could function as a susceptibility gene for ZYMV disease development. On the contrary, ZYMV-CP levels were high in *ClaRBOHB*-silenced watermelon plants (Figure 6B) suggesting that *ClaRBOHB* is probably a gene of resistance against ZYMV. Watermelon plants were observed for 30 days after ZYMV inoculation; at 21 dpi all control and *ClaRBOHB*-silenced plants were infected (100 %) while only 60 % of *ClaATRIP*-silenced plants exhibited ZYMV infection symptoms.

**Fig. 6.**
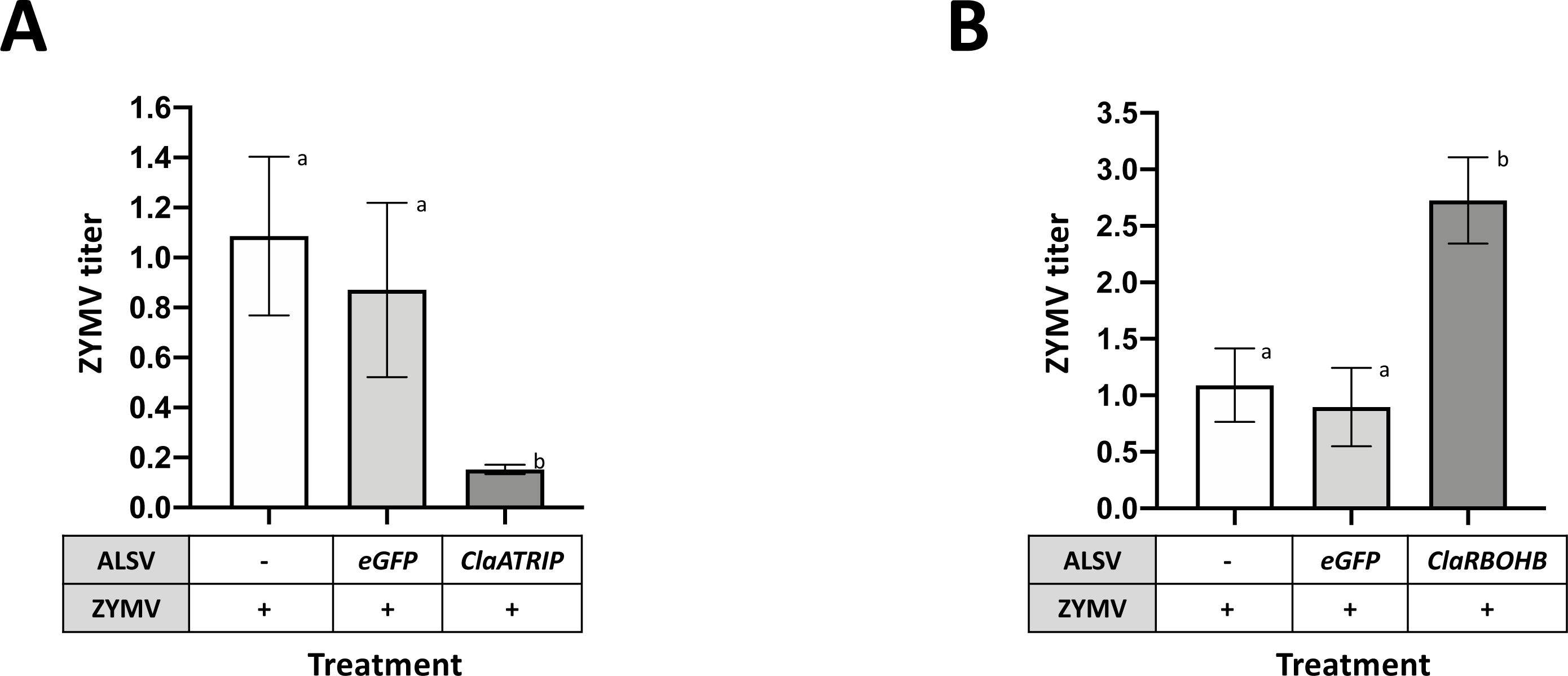
Quantification of ZYMV CP in silenced for A. *ClaATRIP* and B. *ClaRBOHB* watermelons at 15 dpi with ALSV from *N. benthamiana* extract and at 3 dpi with ZYMV inoculation. Quantification analysis performed by Schmittgen-Livak method. Statistical comparison of means done by T-test for significance, p<0.05. White column: ZYMV; grey column: *eGFP*; black column: *ClaATRIP* or *ClaRBOHB*.

## 4. Discussion

In this study we investigated the role of miRNAs in watermelon’s interaction with ZYMV. It is the first time that watermelon miRNAs are registered in the miRBase. Several of the miRNAs found were known (22) but the majority were new ones (331).

Virus infections result in a dramatic transcriptional reprogramming to its host. In the present study of watermelon-ZYMV interaction, 197 miRNAs were downregulated (Figure 2A) and 107 miRNAs were upregulated (Figure 2A) upon ZYMV infection was observed. This may indicate an effort of the plant to increase its transcriptomic activity to establish resistance upon ZYMV infection or the effort of the virus to make the host environment more suitable for its replication. Healthy plants expressed 197 miRNAs, some of them may be regulating post-transcriptionally the expression of resistance factors in order to avoid fitness costs. In particular, there were 34 resistance genes (CC-NBS-LRR, TIR-NBS-LRR, TIR-NBS) predicted as targets of 32 miRNAs (24 miRNAs were uniquely expressed in healthy tissue and silenced upon ZYMV infection) in healthy watermelon. The most abundant size miRNA clusters in watermelon-ZYMV interaction were the 23- and 24-nt-long miRNAs (in both the mock and ZYMV-infected tissue); in contrast watermelon infected by CGMMV resulted in producing high numbers of 23-nt long miRNAs while in the mock the 21- and 22-nt long were predominant. This difference may be due to the fact that ZYMV and CGMMV infecting watermelon belong to potyvirus and tobamovirus plant virus families, having a completely different replication and expression strategy of their genomes *in planta*. The reduced number of miRNAs upon ZYMV infection, observed in our study, is in agreement with results in other plant-virus interactions (e.g. maize-sugarcane mosaic virus (SCMV) [57], watermelon-CGMMV [58], rice-rice stripe virus [59]). The presence of the potyviral silencing suppressor i.e. HC-Pro, with multiple functions such as binding small RNAs [60] and prohibiting HEN1 to methylate small RNAs [61], may play a crucial role reducing the number of miRNAs in infected tissue [62].

Plant miRNAs play a role in defence against virus infection in a process called miRNA-mediated resistance. All the plant miRNAs, available in the miRbase, were tested for targeting the ZYMV genome [53]. The mir396a-3p, miR8706a, and miR1886i-5p (with expectation score of 2.0) could be targeting the sense and reverse complement of the ZYMV genome, respectively; none of them was detected in watermelon miRNAome. All the above miRNAs are predicted to regulate the ZYMV RNA at the cleavage step of the post-transcriptional level. However, for confirmation purposes of the regulation the ZYMV RNA target by these miRNAs and the possible regulation mechanism further experimentation is needed. The watermelon miRNAome were tested for targeting the ZYMV genome. *In silico* detection of watermelon miRNAs able to target the ZYMV genome did not result in such probable miRNAs since their expectation scores were > 3.5. These findings imply that in watermelon there is no direct miRNA-mediated antiviral activity since the direct post-transcriptional regulation of ZYMV RNA does not seem, at least bioinformatically, possible. However, small nucleotidic modification via gene editing could render a miRNA permissive to induce ZYMV silencing (inhibition by mRNA cleavage or by mRNA translation) and this approach requires future testing. Furthermore, the identified miRNAs that regulate watermelon’s resistance/susceptibility genes and/or ZYMV RNA could be evaluated via an amiRNA-mediated approach that has been established in watermelon [36].

MiRNA-mediated resistance in plants could be achieved through the regulation of resistance or susceptibility factors. In particular, de-repression of resistance genes and suppression of susceptibility genes through manipulation of miRNAs could induce the desirable resistance against virus infection. The 34 resistant genes with the respective 32 miRNAs could suggest that with those resistance genes and the identified miRNAs might be involved in ZYMV infection and this requires further studying aiming at inducing genetic resistance against ZYMV in watermelon. The differentially expressed miRNAs in ZYMV-infected watermelon were identified bioinformatically as well as their endogenous target genes, and we validated the differential expression of ten target genes via RT-qPCR upon ZYMV infection, confirming the hypothesis set that ZYMV infection affects the expression of these genes.

In the present work we confirmed that ALSV is an effective vector for VIGS in watermelon although there was a need to have *N. benthamiana* as an intermediate host for ALSV amplification as suggested in [23]. ALSV-infected watermelon plants did not exhibit visible disease symptoms except a shorter stature and we managed to silence successfully three endogenous genes, namely *ClaPDS*, *ClaATRIP*, and *ClaRBOHB*. Τhis is consistent with what has been reported by Tamura *et al*. (2013) [63] and Igarashi *et al*. (2009) [20]. A fragment of 200 bp is sufficient to induce silencing for *ClaATRIP* and *ClaRBOHB*, and GoldenBraid is an easy method for creating the constructs.

We assessed the disease phenotypes for control and VIGS plants and our results suggest that the two selected genes appear to play a role in ZYMV infection of watermelon. In particular, when *ClaATRIP* was silenced, the ZYMV infection was reduced in relation to controls. The pre-inoculation with transformed ALSV::*ClaATRIP* protected watermelon by 40 %. This protection is not due to the *in planta* competition of the two viruses (ALSV and ZYMV) or their synergy, because the ZYMV titer from ALSV::*eGFP*-treated plants was similar to the titer of ZYMV in the absence of ALSV (Figure 6). On the other hand, when *ClaRBOHB* was silenced (loss-of-function) the ZYMV infection was facilitated; at the end of the observation period all watermelon plants were infected. It could be suggested that overexpression of *ClaRBOHB* gene could protect the crop from the ZYMV infection. Based on the data presented here, we could suggest that *ClaATRIP* is a susceptibility gene, while *ClaRBOHB* is a resistance one. Further evidence is needed to confirm these findings via gene editing and/or overexpression of these watermelon genes and subsequent study of the watermelon-ZYMV interaction. Our finding could be used to create “RNA-based vaccines” against *ClaATRIP* in order to protect watermelon against a serious disease as the one from ZYMV.

## 5. Conclusions

This work provided information on the miRNAs of watermelon. In addition, the VIGS technology employing the ALSV vector functioned successfully in watermelon. Lastly, and most importantly, the loss-of-function of *ClaATRIP* and *ClaRBOHB* genes of watermelon indicated that *ClaATRIP* is a susceptibility gene and *ClaRBOHB* a resistance gene against ZYMV infection.

## Statement of author contributions

M. Berbati performed the experiments, analyzed the data, and wrote the draft manuscript; AK assisted in the experiments and the data analysis; M. Bousali and TK performed the miRNA bioinformatics analysis; AV designed the research project and supervised the present research work; TM created the ALSV vector/plasmid constructs. AV, AK, TM, and TK edited the manuscript.

## Funding

This study was supported by a scholarship to M. Berbati by I.KY. (grant 2018-050-0502-13750) and the Ministry of Education, Youth and Sports of Czech Republic (grant COST LTC20066 Genome Editing in Plants) granted to TM.

## Declaration of competing interest

The authors declare that they have no known competing financial interests or personal relationships that could have appeared to influence the work reported in this paper.

## Data availability

Data of miRNAs were deposited in the miRbase. Data will be made available upon request.

## Acknowledgement

Authors wish to thank A.E. Dorrance (The Ohio State University, Wooster, USA) for providing the original ALSV vector. The Agricultural University of Athens is acknowledged for providing space and facilities to carry out this project.

## Compliance with ethical standards

This research did not involve any human and/or animal participants. All authors declare that they have no conflict of interests.

## Notes

### Competing Interest Statement

The authors have declared no competing interest.

