## Supplementary figures and images for "Identification of watermelon genes involved in the ZYMV interaction through a miRNA bioinformatics analysis and characterization of *ATRIP* and *RBOHB*"

### Fig. S1-NbPDS_SILENCING_v3.pdf

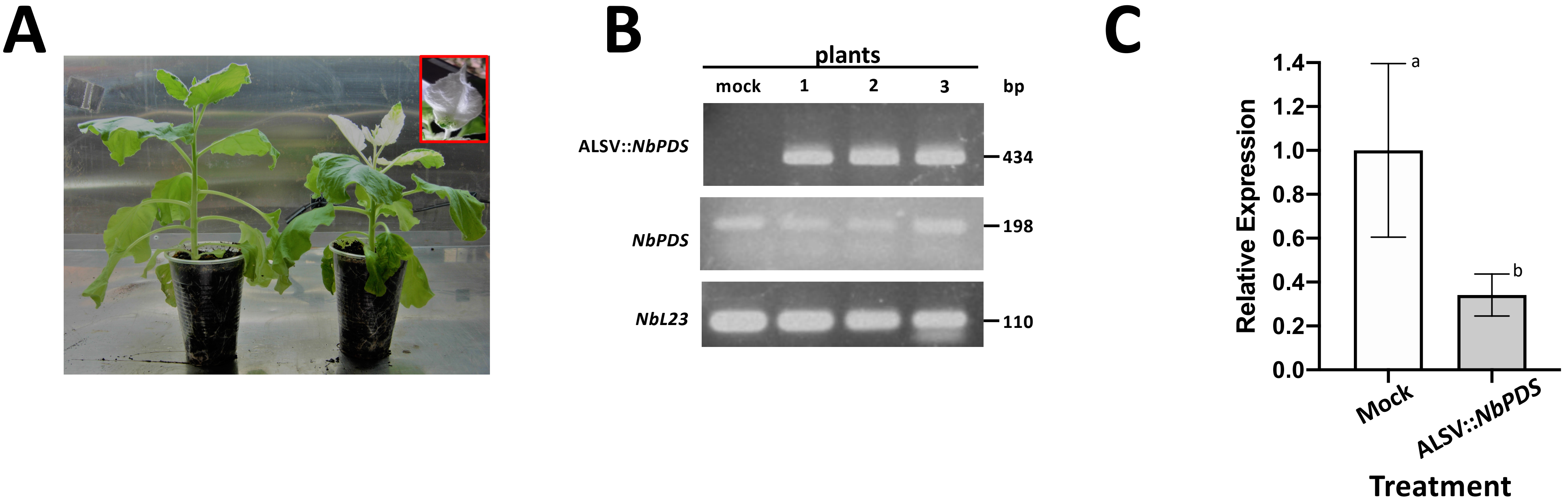
